# Experimental breeding reveals mating patterns and F1 phenotypic diversification in the cosmopolitan diatom *Cylindrotheca*

**DOI:** 10.64898/2026.06.14.731673

**Authors:** Hirono Suzuki, Alexandre Détain, Océane Flet, Taïs Ballanger, Anjana Anilkumar, Noémie Corniaux, Joanna Holm, Coline Donat, Matthew C Posewitz, Chris J Hulatt

## Abstract

Diatom mating activity contributes to their enormous phenotypic and genetic diversity, yet little is known about patterns in diatom reproductive compatibility across genetically diverse strains, nor the effects on offspring phenotypes that may confer adaptive evolution, niche partitioning, or trait improvement. Here a panel of 38 Arctic *Cylindrotheca* sp. isolates were crossed pairwise to detect mating compatibility. Positive mating patterns were identified in multiple clades, including amongst crosses of different parental *rbc*L genotypes. F1 isolated from three different crosses presented phenotypic variation in growth rate, plastid traits, and associated photo-physiological responses to blue and green actinic light. Offspring gliding speed and behaviour also varied, providing insights into complex motility traits that link cell morphology, bioenergetics and sensory adaptation with emergent movement patterns. Exploratory analysis of the F1 trait landscape identified a varaible mixture of individual-level and cross-dependent effects, including substantial variation in growth rate between individuals and strong effects of different crosses on morphology and motility. Experimental diatom breeding may offer a unique strategy to study ocean protist evolution and phenotypic diversification and could complement other biotechnological innovations to enhance cultivation yields and crop resilience in mass cultivation.

## Introduction

Sexual reproduction is widespread among protists and plays a central role in accelerating their adaptive diversification (Coelho & Umen 2021, Speijer et al., 2015). Yet despite their global impact in ocean systems, the reproductive biology of microeukaryotes remains poorly resolved (Krueger-Hadfield 2024, Hofstatter et al., 2018). Development of experimental systems for studying protist sex offers new insights into their ecology and creates opportunities for developing improved strains for biotechnology with higher yields and resilience (LaPanse et al., 2021). Meiotic recombination enables trait improvement programs comparable to crop plant breeding (Goulet et al., 2017, Chepurnov et al., 2008), and crossing transgenic strains can be applied to engineer multiple gene-edited traits into offspring (Kong et al., 2023). Despite such potential, the reproductive biology of single celled organisms is challenging to modulate, and studies that capture full protist breeding cycles are rare.

Diatoms are species rich and globally successful ocean primary producers (Mann & Vanormelingen 2013). Sexual reproduction in diatoms is prolific and thought to play a central role in accelerating their adaptive radiation, phenotypic variation, and their large-scale bloom dynamics (Annunziata et al., 2022, Bilcke et al., 2025, Prigent et al., 2025, Ruggiero et al, 2024). The reproductive cells of diatoms have been observed broadly across many taxa, including centric diatoms that display oogamy and homothallic mating (Moore et al., 2017), and particularly pennate diatoms that are generally isogamous and heterothallic (Montesor et al., 2016). However, we lack a systematic understanding of mate compatibility, reproductive barriers, and hybridization zones in these organisms.

The coastal pennate diatom *Cylindrotheca* offers a promising system for addressing these challenges. *Cylindrotheca* is lightly silicified, transformable (Fischer et al., 2002), grows rapidly, and has a broad geographic dispersal with populations adapted to different environments (Glaser et al., 2020, Stock et al., 2019). Sexual reproduction in a small morphotype of *Cylindrotheca closterium* was initially documented by Vanormelingen et al., (2013), who described the possibility to induce mating between clonal isolates of two different mating types that are activated at smaller cell sizes below a sexualization size threshold (SST). Subsequent studies have demonstrated the coordinated expression of genes underpinning cell size sensing and mating activation amongst selected strains (Belišová et al., 2025). *Cylindrotheca* is thought to use a pheromone signalling system functionally similar to *Seminavis robusta* (Klapper et al., 2021), but the molecular logic that determines mate choice and compatibility across this diverse species complex is not fully understood.

Because there are no complete diatom breeding studies comparable to those of plants or yeast (Anand et al., 2023, Naseeb et al., 2021), our initial questions focused on detecting mating between diverse parental strains and exploring the phenotypic trait variation of the offspring. A few studies have shown potential for diatom outcrossing behaviour, including Castelyn et al., (2009) who identified hybridization between genetically distinguishable *Pseudo-nitzchia*. Likewise de Decker et al., (2018) also demonstrated mating between genetically distinguishable *Seminavis robusta*, indicating potential to introduce novel genetic variation to diatom offspring by combining heterozygous parental haplotypes. We hypothesized that exploring *Cylindrotheca* mating patterns across a wide range of genetic and phenotypic diversity could allow us to initially identify broad scale phylogenetic patterns, constraints and opportunities that could be exploited experimentally.

In principal crossing diatoms should have comparable effects on trait inheritance to other domesticated organisms, but there is little evidence how mating leads to phenotypic variation between offspring. Crop plant breeding programs, by analogy, depend upon high-throughput phenotyping to accelerate selection, and so we identified a key requirement to precisely measure the characteristics of individual diatom cells. Because offspring traits shape the fitness landscape of diatoms in ecological and industrial contexts, our second objective was to isolate F1 from crosses of novel parental strains and to determine key cell phenotypes that form the basis of differentiation, competition and selection, for example in the context of ocean microbial ecology or in a selective breeding program.

## Results

### Mating compatibility patterns across *Cylindrotheca* clades

Thirty-eight strains of *Cylindrotheca* sp. were isolated from coastal fjords in Nordland, Arctic Norway, from water samples collected between 2019 and 2024. Selection of strains was based on morphological identification with light microscopy. Cells of different isolates varied in length from 18.6 μm to 128.3 μm, reflecting both phenotypic differences between strains and the extent to which cells had previously undergone natural cell size reduction. A phylogenetic tree of the 38 strains is illustrated in Figure 1, which identifies multiple different *rbc*L clades, of which clades B, C, D, E, F and G share highest BLAST identity with *Cylindrotheca closterium* (Ehrenberg) Reimann & Lewin. Outermost clade A was determined *Cylindrotheca* sp., including for example *cyr*8 that shares 95% BLAST identity with *Cylindrotheca* sp. IIP03 from Lineage I described in Vanormelingen et al., (2013).

**Figure 1.**
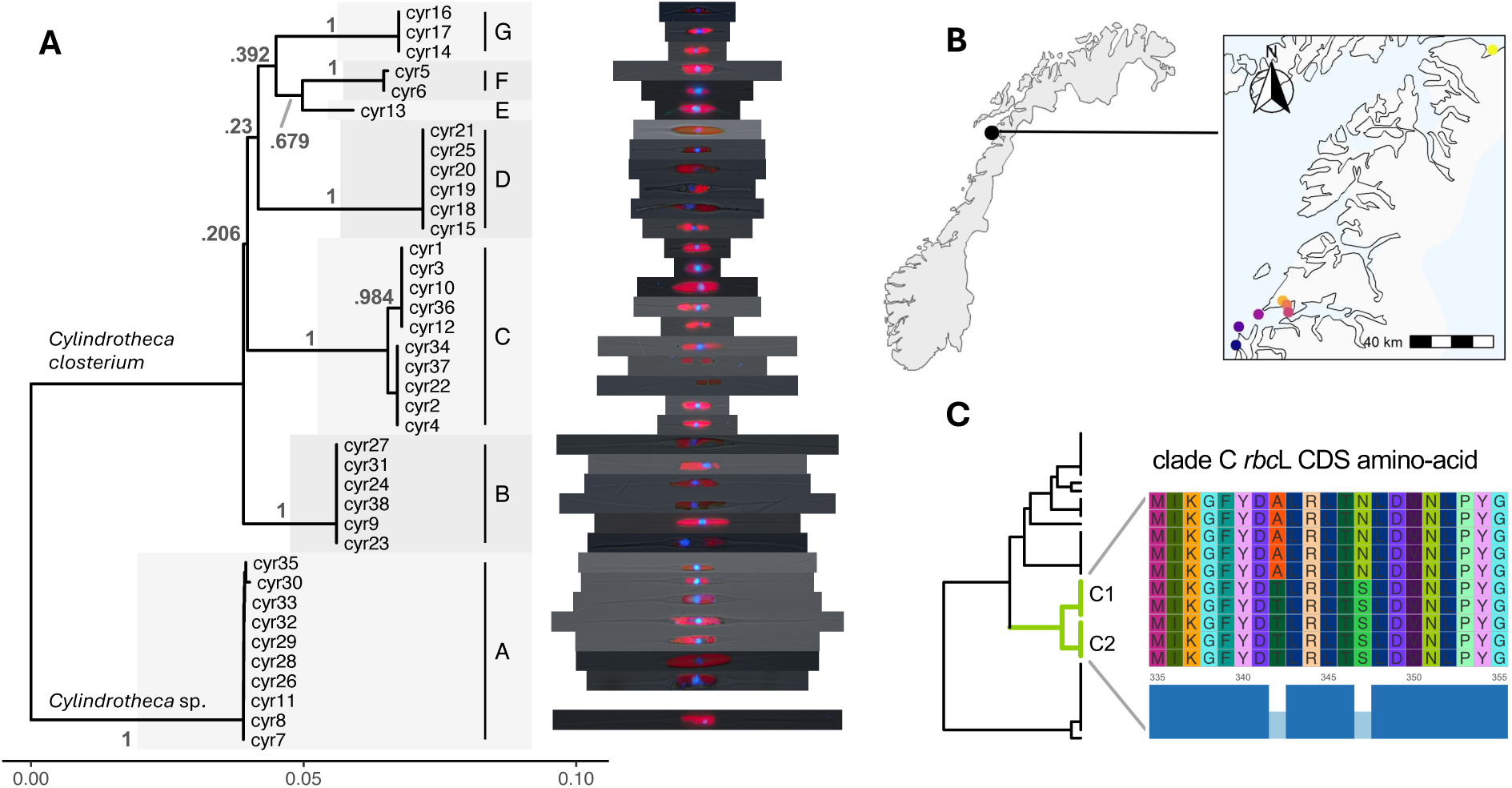
*Cylindrotheca* sp. isolate characterization and phylogeny. (A) Phylogenetic tree of *rbc*L sequences from 38 isolates of *Cylindrotheca* sp. computed with maximum likelihood (ML) and 1,000 bootstrap replicates with the T92 + G model which was selected based on its optimal AICc score. The total alignment length is 1,505 bp and branch support values are displayed. Thumbnails are scaled to relative size, representative images of isolates using combined brightfield, chlorophyll autofluorescence (red) and nuclear labelling (blue) indicating phenotypic variation across clades (*cyr*7 and *cyr*11 not shown). (B) Map of sample site locations along the coastline of Nordland, Arctic Norway. (C) Illustration of the CDS amino-acid sequence variation highlighting variable sites between strains in clade C.

In the first experiment we attempted all pairwise crosses between 38 strains, including controlling for self-mating. Identification of mating activity was defined by observation of the reproductive cells and their progression, including distinctive paired zygotes, auxospores, and large initial cells in each of the replicate wells over 5 days (Figure 2A-M). Expanding auxospores were characterized by the transient appearance of small lipid droplets that coalesced and vanished as cells matured (Figure 2H).

**Figure 2.**
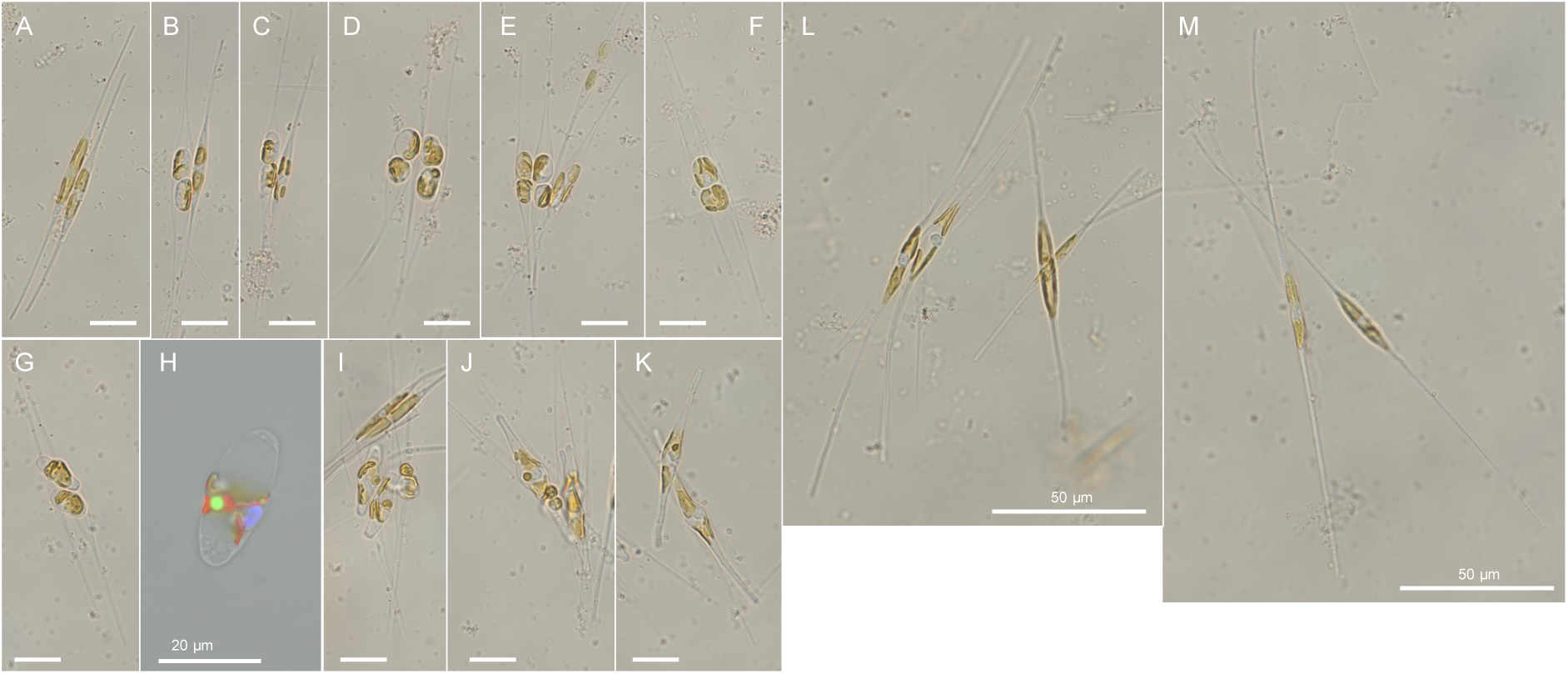
Reproductive cells in a cross between clade B *cyr*24 and *cyr*38 *Cylindrotheca closterium* large morphotype. Progression from paired parental cells through zygotes, to expanding auxospores (A-K) towards initial cell formation (L,M) noting that the different parental cells can be distinguished by their length. It was common in many cases (E,I,J) for more than two cells to be found adjacent. Lipid droplets (H) were a universal, transient and dynamic feature of auxospores. Color in (H) is lipid droplets in green (Bodipy 493/503, blue excitation), chlorophyll autofluorescence in red (blue excitation), and DNA in blue (Hoescht 33342, UV excitation). Brightfield-only images are each proportional in size taken with a 60× objective lens and annotated with a 15 μm scale bar (unmarked, default) or longer 50 μm scale bar (L,M). Combined fluorescence micrograph (H) was obtained with a 100× objective and has an alternative 20 μm size scale bar.

In total we observed mating in 21 crosses between 16 unique strains under minimum *n* = 3 replicated conditions each (Figure 3). Due to the large number of pairwise tests and heterogeneity of activity, mating was scored present or absent. Many crosses however displayed vigorous mating with abundant pairs of sex-derived cells visible in a single microscope field of view (Supplementary data S1). For some crosses only a handful of events were observed in a single plate well, often in localized patches and appearing at variable time frames during the incubation. The overall patterns of mating interactions amongst strains and clades are shown in Figure 3, which is sorted by phylogenetic distance using the tree topology, and includes the cell sizes and growth rates of the isolates. Mating events were observed for *Cylindrotheca closterium* in clades B, C and F, with the highest number of unique crosses observed in clade C. Mating clades B, C and F were reproductively isolated from each other, because inter-clade mating activity was never detected in the test conditions. Clade A included 10 strains of larger and more distantly related *Cylindrotheca* sp. that shared similar or identical *rbc*L haplotypes that did not undergo mating. Clade D, in which 6 isolates were smaller in length, also did not demonstrate any mating activity, whilst clades E and G included few strains.

**Figure 3.**
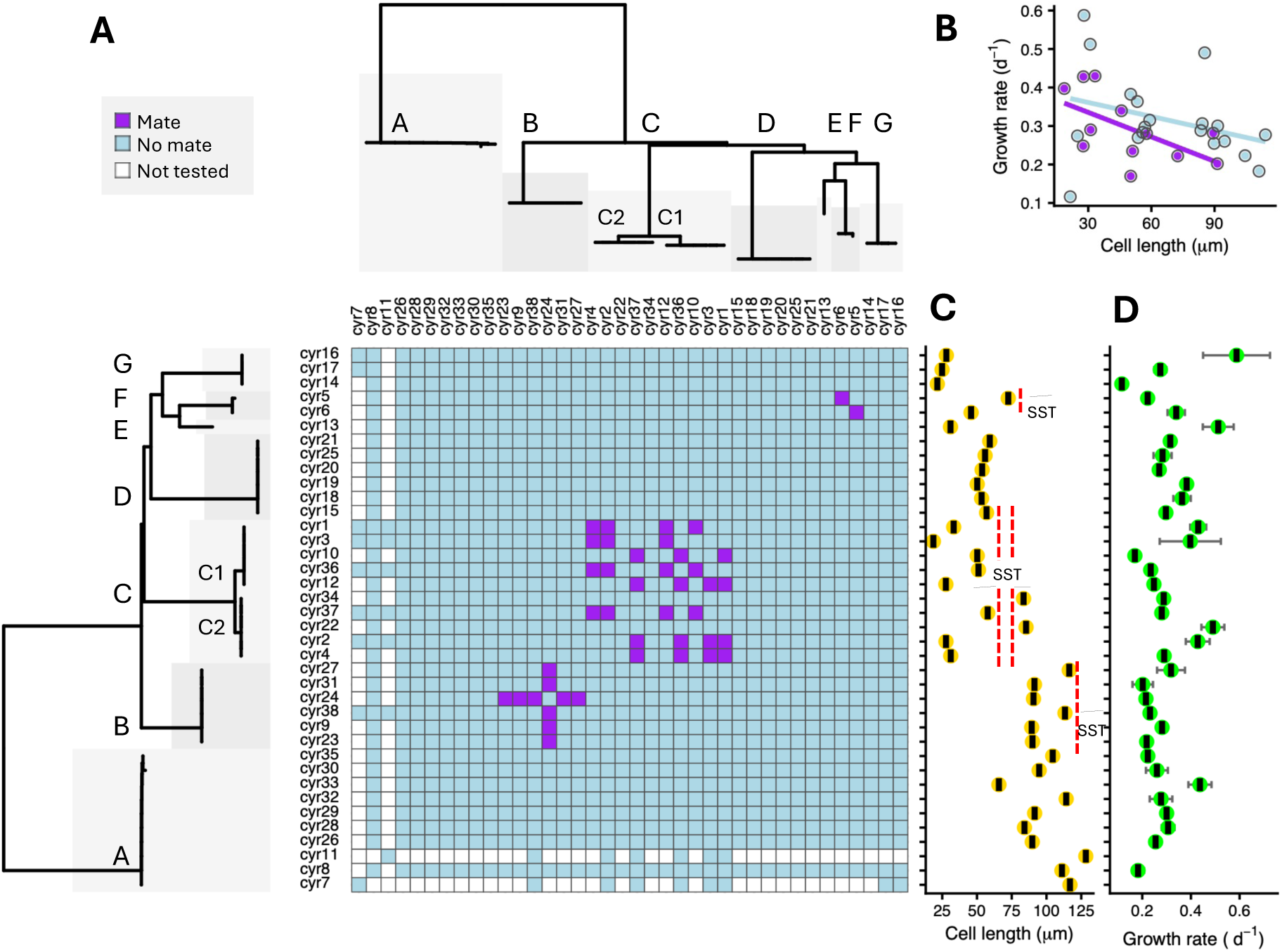
Patterns of mating amongst 38 strains of *Cylindrotheca* sp. Matrix indicating mating and non-mating pairwise combinations of 38 parental *Cylindortheca* isolates which is sorted by phylogeny using the *rbc*L nucleotide sequence tree and annotated with respective clades. The matrix is diagonally symmetrical for visualization of the cell size and growth rate of the parental strains. SST annotation indicates the approximate inferred range of the sexualization size threshold between mating and non-mating isolates in three clades. The correlation between cell length and growth rate of cells that did and did not mate is shown.

Clade C provided 15 mating pairs from 8 of the 10 strains. By examining the mating patterns among strains, we were able to assign two different mating types determined MT1 and MT2 based on rules concordant with heterothallic mating (Table 1). A single exception was the absence of expected mating between strains *cyr3* (MT1) and *cyr10* (MT2), which shared identical *rbc*L haplotypes and each of which mated with two other strains of the opposite mating type. However, at the time of experiments *cyr*3 had become sufficiently small that it was barely motile, which might impact the encounter rate and specific pairwise interactions such that mating was no longer observable in this combination.

**Table 1.** Mating types of *Cylindrotheca closterium* strains from Clade C inferred from replicated crossing experiments (*n*=3) in 24-well plates. Of the 10 strains in Clade C, only the 8 strains actively mating are presented. The assignment of MT1 or MT2 is applied by initially designating strain *cyr*1 as “MT1”. Other strains are referenced relative to strain *cyr*1. Patterns are consistent with two mating types and heterothallic mating compatibility requirements. A single negative result is marked “N” where we expected mating based on the heterothallic mating type rules, but were not able to observe it.

|  |  | Strain | 1 | 2 | 3 | 4 | 10 | 12 | 36 | 37 |
| --- | --- | --- | --- | --- | --- | --- | --- | --- | --- | --- |
|  |  | Mating Type | MT1 | MT2 | MT1 | MT2 | MT2 | MT2 | MT1 | MT1 |
| Strain | Mating Type | Sub-clade | C2 | C1 | C2 | C1 | C2 | C2 | C2 | C1 |
| 1 | MT1 | C2 | × |  |  |  |  |  |  |  |
| 2 | MT2 | C1 | ○ | × |  |  |  |  |  |  |
| 3 | MT1 | C2 | × | ○ | × |  |  |  |  |  |
| 4 | MT2 | C1 | ○ | × | ○ | × |  |  |  |  |
| 10 | MT2 | C2 | ○ | × | N | × | × |  |  |  |
| 12 | MT2 | C2 | ○ | × | ○ | × | × | × |  |  |
| 36 | MT1 | C2 | × | ○ | × | ○ | ○ | ○ | × |  |
| 37 | MT1 | C1 | × | ○ | × | ○ | ○ | ○ | × | × |

Clade C also presented novel mating activity between two genetically distinguished sub-clades designated C1 and C2 that differed in *rbc*L sequence by 6 bp single nucleotide polymorphisms (SNPs), representing the largest 0.4% genetic divergence amongst any of the mating strains in this study (Figure 1A). Translation of the 6 SNPs revealed two AA substitutions in the CDS closely located at positions 342 (Ala/Thr) and 347 (Asn/Ser) in the alignment (Figure 1C). Because clade C included mating cells of each mating type, and additionally two further isolates of larger cells that did not mate, we can infer that the SST for *C. closterium* clade C is approximately between 57.7 μm and 83.5 μm.

*Cylindrotheca closterium* isolates in clade B were much larger than those of clade C, ranging from 89.4 μm to 116.4 μm in length during the crossing experiments. Because all the large clade B cells underwent mating, the SST of *C. closterium* clade B is presumed to be larger than 116.4 μm, approximately twice or more that of clade C, providing evidence of inter-clade variation in SST associated with phenotypic variation in cell size. All of the isolates in clade B shared the same *rbc*L haplotype, and uniquely this appears to be the first observation of mating activity among isolates of the larger *C. closterium* morphotype.

Clade F included only two isolates of *Cylindrotheca closterium* of distinctly unequal size, which nevertheless also underwent mating and implies an SST greater than 72.5 μm. Strain *cyr*5 shared 100% sequence identity with *Cylindrotheca closterium* clade V (NCBI: JX970999) reported by Vanormelingen et al., (2013), and strain *cyr*6 differed from *cyr*5 in 1 bp. Clade F thus represents cosmopolitan *C. closterium* of the smaller morphotype found from the English Channel to Arctic Norway.

Akin with cell size, the growth rate of isolates varied considerably, ranging from 0.12 to 0.59 d^-1^ (Figure 3C, D). Isolates of smaller cells tended to grow more quickly than larger cells, and isolates activated for mating tended to have lower growth rates than non-mating cells (Figure 3B). Notably the two smallest strains, *cyr*3 and *cyr*14, eventually reached the critical survival size threshold (CST) and ceased growth and motility. The large residual variation in the cell size-growth rate relationships thus includes diatom-specific features including SST and CST that influence the overall size-growth patterns across taxa.

### F1 phenotypic variation of controlled crosses

Three sets of controlled crosses were chosen to generate F1 offspring, *cyr*1×*cyr*2 (clade C), *cyr*1×*cyr*4 (clade C) and *cyr*24×*cyr*38 (clade B). Crosses of strains *cyr*1×*cyr*2 and *cyr*1×*cyr*4 were selected with the anticipation that the 6 bp SNPs between their *rbc*L haplotypes was indicative of more extensive genome wide variation and thus presented the best opportunity to produce heterozygous offspring. In addition, we isolated F1 from crosses of *cyr*24×*cyr*38, which had identical *rbc*L haplotypes but represented the larger clade B morphotype, providing novel phenotypic variation amongst *Cylindrotheca* clades. In total 88 F1 offspring were isolated and eventually 20 of these survived to produce viable long-term cultures (Table 2). The F1 cells were each about double the length of the parental cells, including the larger offspring from clade B that measured 175.1 μm compared to 69.8 and 71.4 μm from crosses of the smaller clade C strains.

**Table 2.** Results summary of F1 offspring isolation experiment from three different parental crosses. The clades and genetic distance are indicated, and viability measured by the number of F1 cultures obtained from manually isolated single initial cells. The mean sizes of the parents and F1 offspring were measured using Olympus microscope software. The parents were measured at the time of the mating experiments, and the F1 offspring a few months after establishing the cultures. For each isolate of parental cells 30 cell lengths were measured from the stock cultures. For F1 offspring the mean length of 25 cells from each of three replicate flasks were measured. The F1 length values presented are the overall means of *n* F1 offspring cultures.

| Parental Cross | Clade | Genetic distance *rbcL* SNPs (%) | *rbcL* CDS divergence (AA) | F1 viability | Length parents mating (μm) | Length F1 (μm) |
| --- | --- | --- | --- | --- | --- | --- |
| *cyr1*×*cyr2* | C1 x C2 | 6 bp (0.4%) | 2 | 8/22 (36.4%) | 33.24 × 27.70 | 71.43 (*n*=8) |
| *cyr1*×*cyr4* | C1 x C2 | 6 bp (0.4%) | 2 | 7/39 (17.9%) | 33.24 × 31.12 | 69.75 (*n*=7) |
| *cyr24*×*cyr38* | B x B | 0 bp (0.0%) | 0 | 5/27 (18.5%) | 90.65 × 113.31 | 175.13 (*n*=5) |
|  |  |  | Total | 20/88 (22.7%) |  |  |

The specific growth rate varied ∼3-fold across offspring, with individual F1 from *cyr*1×*cyr*2 and *cyr*1×*cyr*4 presenting the highest growth rates 1.20 ±0.33 d^-1^ and 1.30 ±0.13 d^-1^ respectively (Figure 4A). Noteworthy, several of the individual F1 were visibly different in culture, including isolate “E” from a *cyr*24×*cyr*38 cross, which was paler and grew more slowly than the others, yet consistently propagated at 0.42 ±0.02 d^-1^.

**Figure 4.**
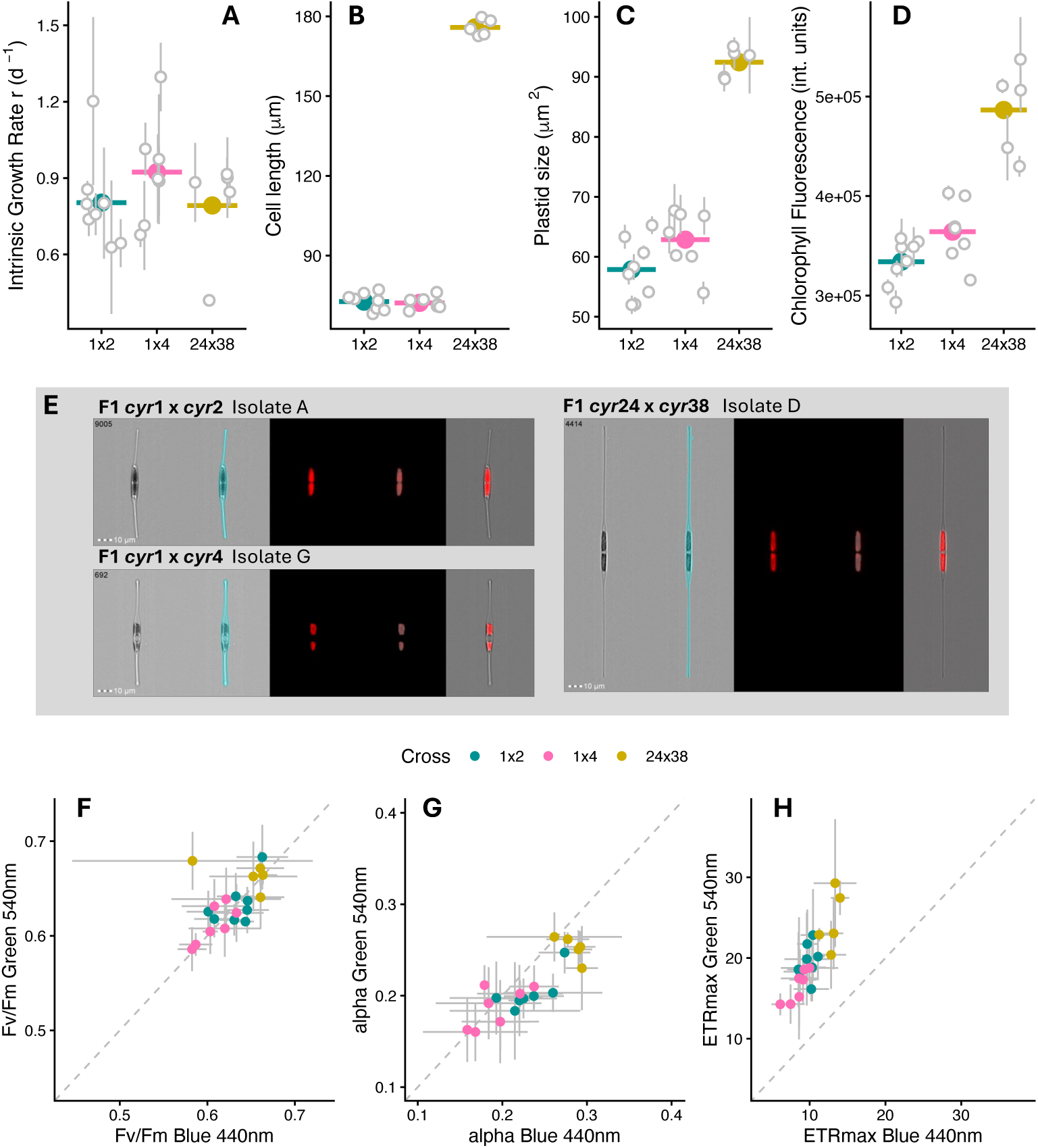
F1 morphologial and photophysiological phenotypes from three parental crosses. (A) Specific growth rate (d^-1^). (B) Cell length (μm) from image analysis (C) Plastid area (μm^2^) from image analysis. (D) Plastid chlorophyll fluorescence intensity (relative units) from image analysis. (E) Example images of individual F1 cells from three different parental crosses and illustration of the image analysis (left to right); Brightfield cell image (Ch4), mask applied to the brightfield image, red chlorophyll fluorescence (Ch5), mask applied to the chlorophyll fluorescence marking the plastid area, composite image of the masked brightfield image and the masked plastid (Ch4 + Ch5 masked). (F) Maximum quantum yield of photosystem II (Fv/Fm) with blue (440 nm) and green (540nm) light. (G) Alpha, α (μmol e- μmol photons^-1^), the initial slope of rapid light curves fitted with the Platt (1980) model under blue (440 nm) and green (540 nm) light. (H) ETRmax (μmol m^-2^ s^-1^ electrons), the maximum photosynthetic electron transport rate of rapid light curves fitted with the Platt (1980) model under blue (440 nm) and green (540 nm) light.

High-throughput single cell imaging cytometry revealed key morphological differences in the cell length, plastid size and chlorophyll content of the F1. Whilst offspring from *cyr*1×*cyr*2 and *cyr*1×*cyr*4 crosses had comparable mean lengths, individual F1 varied from 68.04 ±0.06 μm to 77.15 ±0.01 μm (Figure 4B). The F1 from crosses of *cyr*1×*cyr*4 also tended to have plastid sizes 8.6% larger in area and chlorophyll content 9.1% higher than F1 from *cyr*1×*cyr*2 crosses (Figure 4C, 4D), suggesting parentage effects on bioenergetic organelle features of the smaller clade C morphotype.

Due to the varied plastid traits, we hypothesized that F1 may also exhibit altered photo-physiology. Rapid light curves supplying blue (440 nm) and green (540 nm) actinic light aimed to distinguish whether there was wavelength dependency of plastid functioning. For example, varying amounts and ratios of chlorophyll and fucoxanthin in the fucoxanthin-chlorophyll protein (FCP) complex that might confer altered photosynthetic efficiency or adaptive status. The results show variation in photosystem II quantum yield Fv/Fm, the initial slope α, and the maximum electron transport rate ETR_max_ between crosses and individuals (Figure 4F, 4G, 4H). Offspring of the larger clade B *cyr*24×*cyr*38 crosses were clearly distinguished clustering with elevated Fv/Fm, α, and ETR_max_ relative to F1 from *cyr*1×*cyr*2 and *cyr*1×*cyr*4 crosses. The ETR_max_ of all F1 was substantially green shifted, with offspring showing approximately double the maximum photosynthetic rate under 540 nm actinic light relative to 440 nm. ETR_max_ values of clade B *cyr*24×*cyr*38 offspring were respectively 26% and 49% higher than clade C *cyr*1×*cyr*4 and *cyr*1×*cyr*2 offspring under green light. Similarly, under blue light the ETR_max_ of *cyr*24×*cyr*38 F1 were respectively 31% and 53% higher than *cyr*1×*cyr*4 and *cyr*1×*cyr*2 F1, indicating altered photo-physiological adaptation. Comparable with the other traits, individual offspring also exhibited variation in photosynthetic performance, with more than a 2-fold range in ETR_max_ across all twenty isolates.

Motility phenotypes analyzed by video tracking mapped patterns in gliding speed and behaviour. The mean gliding speed captures the time-integrated movement along tracks, including bursts of movement and stationary periods, whilst the maximum gliding speed represents the highest velocity reached within a track. Patterns show that F1 of the larger clade B morphotypes exhibited faster mean gliding speeds of 4.37 μm s^-1^ and maximum speeds of 11.59 μm s^-1^ compared to their clade C and F counterparts that reached mean speeds of 1.26 and 1.30 μm s^-1^, and maximum speeds of 6.74 and 6.90 μm s^-1^ respectively (Figure 5A). Offspring of *cyr*1×*cyr*2 tended to have ∼14% higher maximum speeds than *cyr*1×*cyr*4 offspring. Behavioural traits also exhibited variation, where F1 from clade C tended to display higher rates of directional change along tracks, with lower confinement ratios than F1 from clade B. Conversely, the greater confinement ratios and lower rates of directional change of the clade B offspring are consistent with higher rates of surface exploration, dispersal and directed gliding migration in the larger *C. closterium* morphotype.

**Figure 5.**
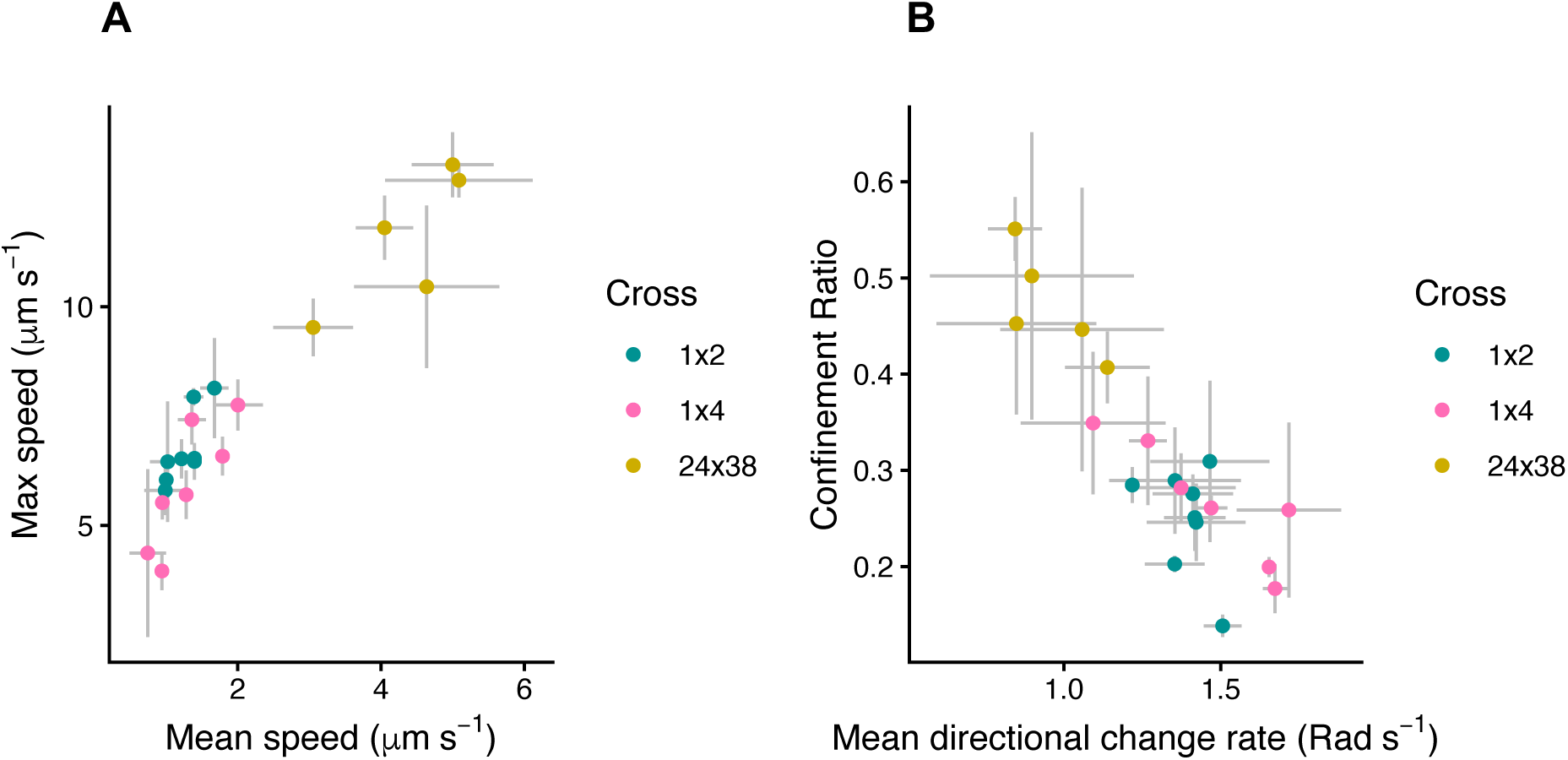
F1 offspring gliding motility phenotypes from three parental crosses. (A) Mean vs maximum cell gliding speed (μm s^-1^) (B) Behavioural traits directional change rate (Radians s^-1^) and the confinement ratio (unitless) that describe spatial dispersal and migration patterns.

To determine which F1 cell features were associated with individual variation vs parentage, mixed-effects models were fitted to 14 different offspring traits (Figure 6A). Random effects models showed that growth rate, which is a master trait derived from interactions among multiple underlying metabolic features, was strongly associated with individual-level variation and parentage effects were negligible. In contrast morphological and motility traits were strongly influenced by parentage, with 47.2-95.2% of variation in these features attributed to the parental cross. PCA analysis (Figure 6B) provides an interpretation the multi-trait landscape of F1 offspring where 83.4% of the total variation was captured by three principal components (Supplementary data S2). The first component (65.5%) could be described as the morphology-motility axis because it featured several traits particularly cell length, plastid area and movement speed with positive contributions. The second component (10.9%) featured primarily macro-metabolic traits including growth and Fv/Fm, whilst the third component (7%) differentiates offspring based on a contrast between growth (−0.63) and electron transport rates, particularly ETRmax under green light (0.43), indicating a trade-off between growth and photosynthetic activity along this axis. Plastid chlorophyll fluorescence also contributed negatively to the third component (−0.39).

**Figure 6.**
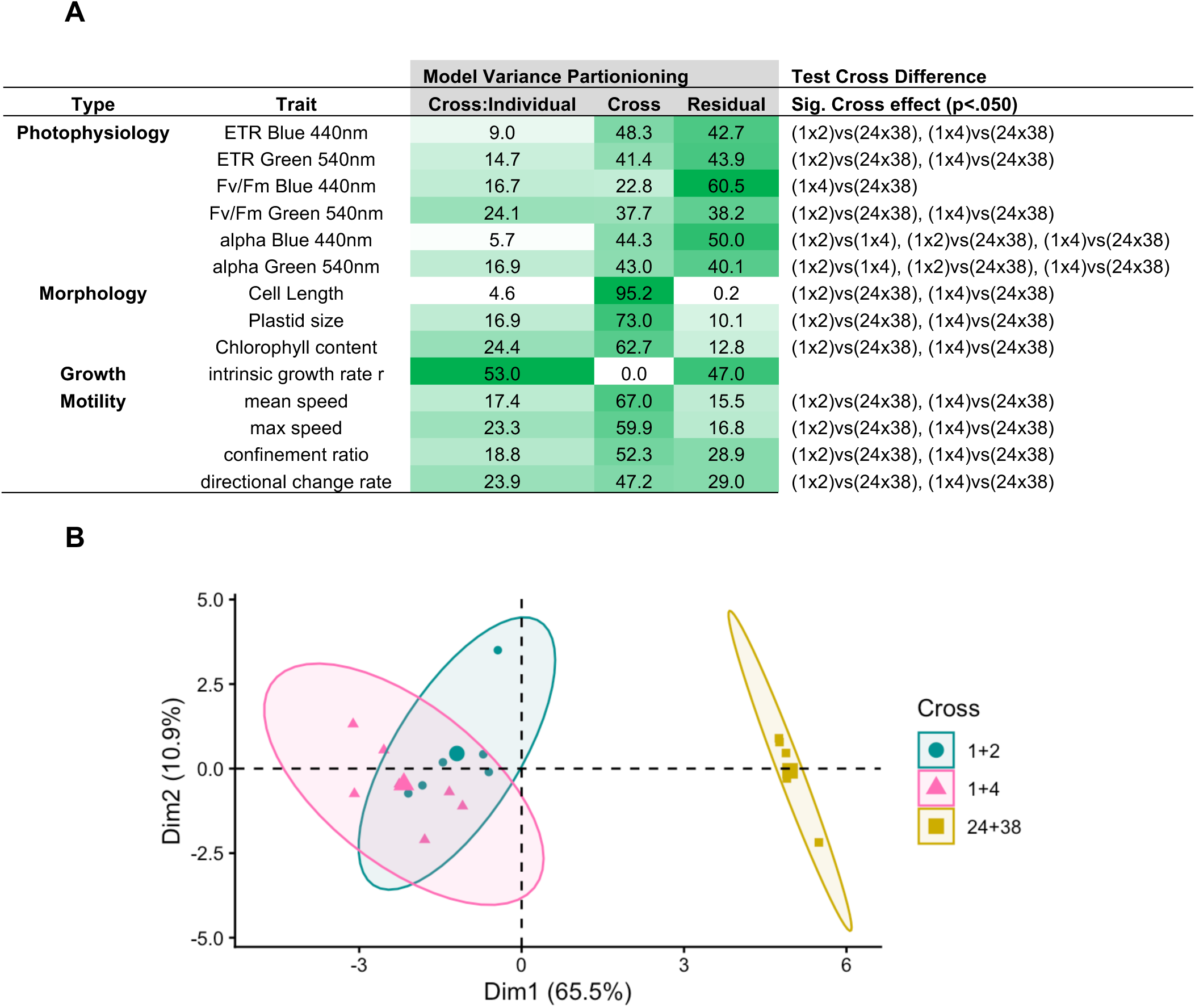
Trait landscapes of *Cylindrothca* F1 offspring. (A) Table summarizing the results of linear mixed effects models for each trait. Models of the from “ *Trait* ∼ 1 + (1 | Cross) + (1 | Cross:Individual) ”, i.e. random effects models, were run for each of the 14 traits and the “Model Variance Partitioning” results represent the percentage (%) of variance explained by individual offspring (Cross: Individual), parentage (Cross) and remaining unexplained variation (Residual). A second model of the form “ *Trait* ∼ Cross + (1 | Cross: Individual) ” was fitted for each of the 14 traits to test whether F1 of different parentage had significantly different traits whilst accounting for individual level variation. Each of the 20 individual F1 in the model included repeated measurements, with *n*=3 (morphology, growth, motility traits) or *n*=4 (photophysiology traits) replicate flasks per isolate. Significant differences between crosses are based on Tukey adjusted p-values presented as “Test Cross Differences” at p<.050. (B) Trait landscape based on PCA analysis of the same 14 traits and three different parental crosses.

## Discussion

Pairwise crosses among diverse *Cylindrotheca* isolates revealed the underlying patterns of diatom reproductive interactions, including mate compatibility and reproductive barriers between clades, that shape population structure, molecular evolution, and emergent evolutionary trajectories. Controlled crosses between different parental strains generated F1 phenotypic variation in growth rate, morphology, photo-physiology and motility traits. Variation in morphology and motility was largely associated with different parental crosses, yet weak parentage effects on growth rate indicate that offspring traits are shaped by varied combinations of parental and individual-level processes that take place during sexual reproduction. Together these findings provide an initial perspective on the fitness landscape generated by diatom sexual reproduction and a framework for exploring the phenotypic diversification, ecological adaptation and applied trait improvement.

### Mating patterns – locating reproductive boundaries and hybrid zones

A key part of the biological species concept is the ability to interbreed (Monnet et al., 2025). Diatoms commonly form extensive species complexes comprising clades of strains with similar appearance and reticulate ancestry, separated by variable genetic distance (Amato et al., 2007, Çiftçi et al., 2022, Mann 1999). Here clades B, C and F of mating *Cylindrotheca* were reproductively isolated from each other, yet multiple strains in subclade C1 mated actively with genetically distinguishable strains in C2 to produce offspring. Hybrid zones, the phylogenetic regions where more distantly related strains remain mate-compatible, are difficult to predict in diatoms, and the underlying molecular mechanisms that determine them are not understood. Nevertheless, heterosis and introgression, for example, could each play key roles in altering diatom fitness and conferring rapid adaptation to novel environments. These are exploitable in a breeding program, but exploration of species barriers and hybrid zones requires targeting parental strain combinations that ideally span broad phylogenetic gradients. Future exploration of species barriers and hybrid zones could be accelerated through high-throughput approaches for targeted parental strain isolation and compatibility screening, including clade-specific molecular markers and fluorescence-based cell sorting of environmental populations.

### Size-growth patterns of *Cylindrotheca*

Diatoms show substantial variation in cell size and morphology that as master traits impact their physiology (Marshall et al., 2012). Broadly across protists, smaller cells tend to grow faster than larger cells, although the pattern is noisy due to altered cell plans and adaptive processes across taxa that compensate for size scaling effects (Kremer et al., 2017). Our smaller *Cylindrotheca* isolates tended to grow more quickly than larger ones, and the isolates that were activated for mating grew more slowly. This is consistent with a previous study showing that sexual reproduction in diatoms is associated with a temporary growth arrest and downregulation of metabolic processes, including photosynthesis, even under nutrient-replete conditions (Annunziata et al., 2022). Such physiological changes are thought to reflect a reallocation of cellular resources toward gamete formation and zygote development, potentially prioritizing reproductive success over vegetative growth.

Overall, clades of smaller *Cylindrotheca* strains probably offer higher growth rates (Figure 3B), which is supported by the enhanced growth of individual clade C offspring (Figure 4A). Although we did not measure the motility of the parental strains, we also note that smaller isolates *cyr*3 and *cyr*14 that reached the critical cell size survival threshold showed little movement. Since the movement patterns of pennate diatoms are central to their ecology and appear to be related to the raphe morphology (Golfier et al., 2026, Bondoc-Naumovitz et al., 2025), cell size reduction is probably linked with movement-associated biotic interactions, including mating, as well as growth.

### Do reproductively isolated *Cylindrotheca* interact?

Marine biofilms are shaped by cellular interactions that structure species abundance-distribution patterns. Raphid pennate diatoms, that glide actively in response to physical and chemical gradients (Apoya-Horton et al., 2006, Zhang et al., 2025), appear to share common pheromone signalling processes involved in mate finding and cell cycle synchronization that are presumably conserved across taxa (Klapper et al., 2021). In *Seminavis robusta*, l-diproline directs gliding behaviour of potential mates, and a pair of SIP+/SIP- pheromones activate cell cycle processes in paired cells of opposite mating types. Despite the clear reproductive isolation observed between clades in this study, it remains possible that inter-clade interactions persist at other biological levels. Our data thus raises the question of whether biotic interactions including cell signalling and gliding movement take place across different clades of *Cylindrotheca*, even in the absence of mating. Diatom decision making (Bondoc et al., 2019), including their receptor mechanisms and molecular logic that guide motility, are not well characterized yet could have important consequences in structuring aquatic microbiomes. Because partner searching and mating has a major fitness cost (Annunziata et al., 2022), pairing decisions and commitment to meiosis are presumably under considerable selection and potentially linked with the rapid emergence of reproductive barriers within diatom species complexes.

### Outcrossing, hybridization and inheritance of cytoplasm

In their study on natural hybridization, Castelyn et al., (2009) reported a genetic distance of 4 synonymous *rbc*L SNPs between mating *Pesudo-nitzchia* sp. strains, which compares with the 6 SNPs and 2 AA acid *rbc*L marker variation between clade C1 and C2 detected in this *Cylindrotheca* study. In principle the AA discrepancy between clade C1 and C2 parents could impact catalytic efficiency or subunit molecular interactions and function of a key protein. However *Cylindrotheca*, like other Phaeophyceae, encodes both the *rbc*L and *rbc*S subunits adjacent to each other in the plastid genome and transcribes them in tandem. Hypothetically the single operon might allow for subunits to coevolve independently in the plastid genome, without functional effects created by altered interactions between nuclear and plastid encoded genes. Thus, in addition to promoting organelle “portability” in a broad evolutionary sense (Grzebyk et al., 2003), such features may also help retain strain compatibility in isogamous sex systems and leads more widely to the question of how diverse protist reproductive strategies interact with and shape their micro- and macroevolution (Collier et al., 2025).

Inheritance of cytoplasm, including mitochondria, plastids, and their genomes, is in many organisms maternal (Yadav et al., 2025, Camus et al., 2022, Chung et al., 2023), but a study on *Pseudo-nitzchia delicatissima* indicates that this is probably not the case amongst pennate diatoms (Ghiron et al., 2007). *Pseudo-nitzchia delicatissima*, which has comparable reproductive dynamics to *Cylindrotheca*, exhibits biparental inheritance of the plastid, where F1 receive and maintain assorted organelles from either parent (D’Alelio & Ruggiero 2015). In plants, studies on biparental inheritance patterns indicate that nucleocytoplasmic interaction and competition between plastids and other cytoplasmic features takes place in hybrid offspring leading to altered metabolic phenotypes, fitness, and survival (Sobanski et al., 2019). With some preliminary high throughput sequence data from 4 individual F1 from *cyr*1×*cyr*2 and *cyr*1×*cyr*4 crosses, we identified representatives of all three parental plastid haplotypes in the offspring, indicating biparental plastid inheritance in *Cylindrotheca*, but no support yet for heteroplasmy (Supplementary data S1). We also detected variation in the size, chlorophyll fluorescence and photo-physiological responses of the plastid to green and blue light between parental crosses and amongst individual offspring. Hypothetically, the inheritance of diatom cytoplasmic compartments (plastids, mitochondria) in combination with diploid (Oliver et al., 2021), allodiploid (Tanaka et al., 2015) or polyploid (Li et al., 2025) nuclear genomes that encode the majority of organelle-targeted genes, might lead to altered F1 fitness in terms of organelle bioenergetics (photosynthesis and respiration), signalling, and related metabolic traits (Chung et al., 2025). Because nuclear-cytoplasmic interactions may shape the fitness and adaptive potential of populations (Ferderer et al., 2025), they may in principle also provide an opportunity for trait optimization in diatom breeding and genetic engineering too.

### Diatom F1 have different gliding speed and behaviour patterns

Gliding provides benthic diatoms with a unique ability to access light, nutrients, and to navigate mate selection in highly dynamic 3D environments, leading to emergent behaviours such as decision-making (Bondoc et al., 2019). Bondoc-Naumovitz et al., (2025) described that different species have widely varying gliding characteristics linked to cell morphology, thus we anticipated that morphologically diverse parents and offspring might show altered motility phenotypes too. Indeed the *Cylindrotheca closterium* offspring displayed substantial variation in motility traits, both between parental crosses and between individual F1. Offspring of the larger clade B morphotype travelled at average and maximum burst speeds ∼3-fold faster than those of the smaller clade C offspring. Additionally, the variation in confinement ratios and directional change rate is associated with altered rates of spatial dispersal that may shape community structure and bloom formation (Détain et al., 2025, Zheng et al., 2023). Because diatom gliding is driven by ATP-dependent actin-myosin motors and cells are guided by environmental gradients (Zhang et al., 2025), motility traits link cell energy metabolism to wider sensing, behaviour and biotic interactions. The implications include for example that larger diatom cells could trade possible growth rate limitations to extract a speed and dispersal advantage that enable them to better compete for resources in fluctuating coastal environments. More widely, the results imply that pennate diatom parentage, competition and selection on offspring gliding phenotypes offers a powerful method of adaptation and cellular reprogramming related to complex behaviour and individual fitness.

### Innovation and applied aspects

Crossing experiments are fast when strain numbers are low, but the all-pairwise approach is impractical to scale. A wider breeding program requires optimization of the search space for a large number of prospective parents. For example, designing a sparse matrix or selecting crosses that traverse specific phylogenetic regions of interest. With some initial data it is possible to substantially increase search efficiency. Simple PCR barcodes, cell length and known mating type can quickly guide mate selection, eliminate unlikely combinations, and substantially increase efficiency in hypothesis testing (Table 3).

**Table 3.** Summary of the challenges and innovation solutions identified for implementing a scalable diatom breeding program.

| Breeding component | Problem or need | Potential solutions |
| --- | --- | --- |
| Parental strain isolation and crossing. | Increase phenotypic and genetic diversity of compatible parental strains.<br>Predict mating compatibility to optimize the mating search space. | High throughput isolation e.g. cell sorting, selection media, to increase the number of strains.<br>Use cell image features, length, and PCR barcodes to index strains likely to be compatible.<br>Once mating is known, apply heterothallic mating-type rules to reduce the search space. |
| Isolation of F1 offspring | Isolation of the offspring with pipette is time consuming, low throughput, and has a moderate success rate. | Imaging, robotics, and automation for offspring cell selection.<br>Selection of mating surfaces and treatments to alter adherence and retention of mating cells.<br>Selectable markers for double transformants. |
| Phenotyping F1 | Increase throughput, sensitivity and range of traits. | Design high throughput phenotyping and selection arrays for traits such as growth rate across light and thermal gradients.<br>Flow cytometry probes for high throughput image quantification of cell features. |
| Maintenance of strains | Time investment to maintain large numbers of parents and F1 offspring for experiments is high.<br>Strains become non-viable below a critical size threshold. | Robotics and automation<br>Cryopreservation, minimize growth rate in culture. |
| F2 generation and backcrossing | Need to accelerate cell size reduction to activate mating in F1.<br>In plant breeding, backcrossing with parental lineages is common, but in diatoms mating activation of F1 and parental lineages is temporally separated. | Rapid cultivation of F1 to reduce cell size, microsurgery (Belišová et al., 2025) to activate mating.<br>Cryopreservation of parental strains to delay cell size reduction. |

Because diatom breeding is labour intensive, robotics and automation offer natural opportunities toward increasing throughput (Huang et al., 2023). A key bottleneck is isolation of the offspring. To our knowledge all studies have achieved this by hand pipetting, which remains difficult to scale because it is time consuming, the cells are fragile, and F1 growth initially lags during which time parental strains overgrow them. Comparatively in the chorophyte *Chlamydomonas*, for example, the offspring selection problem has been routinely solved by exposing mating cells to chloroform to select for the more robust zygotes. Clearly other methods are needed for diatoms, potentially including the introduction of selectable markers into parental cells and isolation of double-positive offspring, or the use of automated cell picking (Table 3).

Breeding programs typically require multiple generations of selection, but diatoms experience a protracted interval between establishment of the F1 and activation of mating to produce F2. The ability to reduce the interval from initial cell to SST activation e.g. via manual shortening (Belišová et al., 2025) is not only important for throughput but required to overlap the mating window of F1 with older parental lineages, for example to achieve backcrossing. Techniques such as cryopreservation to stall the size reduction of the parents, combined with accelerated growth or shortening of the F1 cells, could offer feasible solutions to partially synchronize the reproductive windows of successive generations (Table 3).

Diatoms used in mass cultivation systems today are often wild-type strains isolated years or decades ago, and variations in yield are common (LaPanse et al., 2023). The ability to induce and control mating in a diatom “hatchery” environment offers advantages for maintaining cultivars at peak fitness, by supplying a stream of new F1 for scaleup. Because our results indicate large variation in F1 growth rates and photo-physiological traits, gains in raceway pond production could potentially be obtained by crossing and selecting diatoms and could compound benefits of complimentary strategies including random mutagenesis (Jankowicz-Cieslak et al., 2016, Arora & Philippidis 2021), gene editing (Krishnan et al., 2023) and experimental evolution (LaPanse et al., 2021) for increasing yield and environmental resilience. Such complimentary approaches could generate novel phenotypic variation that may subsequently be combined and stabilized through sexual crossing and selection, expanding the accessible breeding landscape beyond naturally occurring variation.

### Materials & Methods Strain isolation

Thirty-eight strains of *Cylindrotheca* sp. were isolated from multiple coastal sites near Bodø in Arctic Norway between 2019 and 2024 (Figure 1) using enrichment cultures, dilution in multi-well plates and manual pipetting on an Olympus CKX53 inverted microscope (Olympus Europa GmbH, Hamburg, Germany). Cultures were maintained at 15 or 16°C under low light and pre-cultured in 50 mL flasks containing f/2 medium (Guillard & Ryther 1962). Strains were included in mating experiments based on their broad morphologic identity to the genus *Cylindrotheca*, without other selection criteria. The initial crossing experiment was blinded to the strain sequence identity, because we constructed the *rbc*L phylogeny afterwards.

### Mating experiments

Crossing experiments were carried out in 24-well plates in f/2 medium as described by Vanormelingen et al., (2013). The temperature was 16°C with light intensity approximately 7-8 μmol m^-2^ s^-1^ photosynthetically active radiation (PAR) measured with a Li-Cor LI-190 sensor. Mating activity was examined daily with an Olympus CKX53 inverted microscope over a duration of 5 days, to ensure we were able to detect cell progression in delayed onset or low-abundance mating. Where mating was not observed, infrequent, or heterogenous, we attempted to exhaustively scan the entire well, whilst crosses with positive, vigorous mating, we examined at least the first 20-100 or more events. Concordance of at least three replicate wells confirmed a positive mating result.

The growth rate of isolates was established by culturing each strain in triplicate 250 mL flasks containing 50 mL f/2 medium in an incubator at 15°C under 7 μmol m^-2^ s^-1^ PAR. The optical density (750 nm) of resuspended cells was measured daily and the growth rate (μ, d^-1^) was estimated with equation (1).

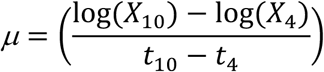

Where *X* is the optical density at day 10 (*X*_10_) and day 4 (*X*_4_), and *t* is the time (d) at day 4 (*t*_4_) and day 10 (*t*_10_).

The cell length of each of the parental strains at the time of mating experiments was measured with an Olympus BX43 microscope with an Olympus DP28 camera and CellSens v.3.2 software with a ×40 objective.

### DNA extraction and sequencing

DNA was extracted from pelleted cells using either an E.Z.N.A.® HP Plant DNA Kit (Omega Bio-tek, Georgia, USA), or cells were directly squashed on microscope slides. For DNA extraction from the cell pellets, bead milling (5,000 rpm for 30s, Precellys evolution homogenizer, Bertin Technologies, Montigny-le-Bretonneux, France) with 0.1 mm glass beads were used for cell lysis. DNA quality and quantity was checked with a NanoDrop ND-1000 (Thermo Scientific, MI, USA). Where only a low amount of cells were available, a few hundred cells were transferred to a microscope slide and burst by sliding the sterile pipette tips onto the cells after allowing it to dry at ambient temperature. The pipette tips and slide were immediately washed with 8 μL of molecular analysis grade water and collected into PCR tubes. PCR amplification of the *rbc*L was conducted using the extracted DNA. PCR amplification was performed using the forward primer DPrbcL1 (Daugbjerg and Andersen, 1997) and reverse primer DPrbcL7 (Daugbjerg and Andersen, 1997) with a total volume of 20μL, consisting of 10μL Phire green hot start II PCR Master Mix (Thermo Scientific, Lithuania), 1μL of each respective foward and reverse primer, 7μL of water, and 1μL of extracted DNA. We used a initial denaturation at 98°C for 5s, followed by 29 cycles of 98°C for 5s, 47.1°C for 30s, 72°C for 28s, with final extension at 72°C for 1 min. Amplified PCR products were separated and visualized by electrophoresis in 1.2 % agar gel using SYBR safe dye (Thermo Scientific, Vann allen way, Canada), and a 1kb ladder (Invitrogen, Carlsbad, CA, USA) to check fragment sizes. PCR products with desired size were purified and sequenced by Macrogen (Amsterdam, Netherlands) using primers NDrbcL12R (Daugbjerg and Andersen, 1997), NDrbcL13F (Daugbjerg and Andersen, 1997), NDrbcL6F (Daugbjerg and Andersen, 1997), 17R (Jones et al., 2005), and 15R (Jones et al., 2005). These sequencing primers in combination yielded five overlapping sequence reads per PCR reaction, and after manual curation and trimming for sequence quality were assembled into high confidence *rbc*L sequences using their encoded base quality scores of the .ab1 files in Geneious Prime (2025.0.3).

### Phylogeny

The *rbc*L sequences were aligned with MUSCLE, manually inspected, and the ends trimmed to a total alignment length of 1,491 bp. The phylogeny was constructed in MEGA11. First the optimal model was identified as the T92+G model based on comparison of AICc scores. The optimal tree was then computed using Maximum Likelihood and 1,000 bootstrap replicates. To examine the protein sequence, the reading frame of the curated *rbc*L sequence alignments were annotated in Geneious Prime allowing vizualization of the AA sequence alignment. The sequences of the 38 strains are available in NCBI GenBank accession numbers PZ322834-PZ322871 and in the Zenodo repository as a .fasta encoded nucleotide alignment file, a Newick tree built with the nucleotide alignment and the optimal model in MEGA (Figures 1 and 3), and a .fasta encoded amino-acid alignment of the CDS.

### Isolation of F1 offspring

F1 offspring were obtained by conducting mass crossing experiments in well plates and 10 cm glass petri dishes. Initial cells were manually isolated using a 10 μL pipette and placed as single cells into 16 mm glass tubes in f/2 medium and incubated at 15°C under low light <5 μmol m^-2^ s^-1^ PAR. Growing cultures could be observed in the tubes after 6 to 8 weeks, at which point a small sample of each was inspected under the microscope to confirm the cultures were the larger offspring and were not contaminated by the parental cells.

### F1 offspring phenotypes

Twenty F1 isolates were each cultivated in triplicate in 16 mL of f/2 medium in 100 mL flasks, which were placed in an incubator at 15°C at 7-8 μmol m^-2^ s^-1^ PAR. The resuspended cell density of each flask was measured daily by haemocytometer cell counts. Cells were counted daily on five 1 mm^2^ grids and the mean cell density (cells mL^-1^) was calculated. The intrinsic growth rates were modelled with the logistic equation (2).

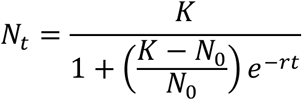

Where *N*_0_ is the initial cell density, *N_t_* is the cell density at time *t*, *K* is the maximum cell density, the upper asymptote, and *r* is the intrinsic growth rate (d^-1^).

The length of the F1 cells was measured by microscope with the identical method to the parental cells, so that the sizes of parents and offspring could be directly compared in Table 2. In addition, the F1 cell size was secondly measured with high throughput imaging flow cytometry, amongst the other morphological traits. Imaging flow cytometry (IFC) was conducted with an Amnis ImageStream equipped with a ×40 objective, a 488 nm blue laser operated at 0.5 mW, and a 785 nm laser operated at 5.0 mW. Image data was collected on three channels: Brightfield (BF) on Ch4 (595-642 nm), chlorophyll fluorescence on Ch5 (642-745 nm), and long-wavelength side scatter (745-780 nm) was used to positively detect the outline of the long transparent apices of the cells during cell image acquisition. Image analysis was conducted with IDEAS v.6.2 software using fluorescence compensation and applying a template to all samples. The cell and plastid morphology were detected using image masking with the “adaptive erode” and “morphology” functions. Cell features including the length (Ch4 masked length, μm), plastid size (Ch5 masked image area μm^2^) and chlorophyll intensity (Ch5 masked image intensity units) were extracted.

For microscope fluorescence imaging cells were labelled with Hoechst 33342 (Thermo Fisher Scientific, Germany) at a final concentration of 1.0 μg ml⁻¹ and BODIPY 493/503 (Thermo Fisher Scientific, USA) at 100 μg ml⁻¹ and incubated for 10 min in the dark. Stained cells were observed by epifluorescence microscopy using UV LED excitation light for Hoechst and blue LED excitation for BODIPY and chlorophyll fluorescence.

### Photo-physiology

Photosynthetic traits were measured with a WALZ Multi-Color PAM (Heinz Walz GmbH, Effeltrich, Germany). Cells were dark-adapted for 20 minutes at 15°C then rapid light curves were made recording initially *Fv*/*Fm* in the dark-adapted condition followed by Y(II) under increasing actinic light intensity. Both blue (440 nm) and green (540 nm) actinic light were used in separate recordings with a 30 s actinic light step duration. A Li-Cor US-SQS/L miniature PAR sensor was placed inside the quartz cuvette to record the in-situ actinic irradiance in each sample. The electron transport rate (ETR, μmol m^-2^ s^-1^ electrons) was determined in Walz PamWin3 software (3).

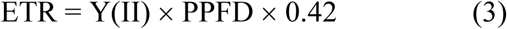

Where Y(II) is the quantum yield under illumination by actinic light i.e. Δ*F*/*F_m_*′, PPFD is the intensity of either blue (440 nm) or green (540 nm) actinic light (μmol m^-2^ s^-1^ PAR) and 0.42 is a standard coefficient of light absorption by photosystem II. Rapid light curves were fitted using the model of Platt et al. (1980) (4):

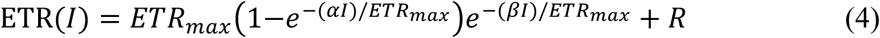

Where ETR(*I*) is the electron transport rate at irradiance *I*, ETR_max_ is the maximum electron transport rate (μmol m^-2^ s^-1^), α is the initial slope of the light response curve i.e. the photosynthetic efficiency; μmol e- μmol photons^-1^, and *β* describes photoinhibition at high irradiance. The parameter *R* represents a baseline offset corresponding to the y-intercept i.e. ETR at zero irradiance. For each of the 20 offspring, 4 replicate flasks were measured with blue (440 nm) and green (540 nm) actinic light.

### Motility

Gliding motility was analyzed by video microscopy using an Olympus BX43 microscope fitted with a DP28 camera and ×10 objective. Cells were pipetted into 20 µm depth sperm counting chamber slides to allow free movement (CV 1020-2cv, CellVision technologies, The Netherlands). Videos of 90 s were recorded at a resolution of 1080p at 1fps in .avi format. Recorded videos were then converted into black and white image sequences with a resolution of 720p using a python script. Using the Trackmate plugin for Fiji (imageJ), image sequences were loaded for tracking cell particles and acquiring movement parameters. Difference of Gaussian (DoG) was used to detect particles with an object diameter of 20 pixels, initial quality thresholding at 0.68 and quality filter on spots set at 0.97. The Linear Assignment Problem mathematical framework (LAP tracker) was used for tracking particles with a frame to frame linking of 30 pixels and gap closing of maximum 50 pixels and 30 frames. Tracks under 23/90 frames were removed.

### Analysis & statistics

Data was analyzed with R. Mixed effects models used the package “lme4” and evaluation of the fixed parental effects using “emmeans”. The rapid light curve parameters were modelled using the “platt” package. Offspring growth rates were modelled with the “growthcurver” package.

### Data Availability

Sequence data are available at NCBI under GenBank accession numbers PZ322834-PZ322871. Data in the manuscript are available at Zenodo: https://doi.org/10.5281/zenodo.18982249 *<DATA will be released on publication and are made available by private link for peer review>*

### Supplementary data

A single Supplementary data file is associated with the manuscript.

## Author Contributions

CJH conceptualized the research project with MP. HS and CJH conducted strain isolation, crossing experiments and molecular identification of parental strains with support from NC. HS conducted offspring isolation. CJH conducted F1 phenotyping with support from AD, OF, TB and JH. AA and CD provided input to the original draft and all authors approve the final manuscript.

## References

1. Amato, A., Kooistra, W.H.C., Ghiron, J.H.L., Mann, D.G., Pröschold, T. & Montresor, M. (2007). Reproductive isolation among sympatric cryptic species in marine diatoms. Protist 158, 193–207.

2. Anand, A., Subramanian, M. & Kar, D. (2023). Breeding techniques to dispense higher genetic gains. Frontiers in Plant Science 13, 1076094.

3. Annunziata, R., Mele, B.H., Marotta, P., Volpe, M., Entrambasaguas, L., Mager, S., Stec, K., d’Alcalà, M.R., Sanges, R., Finazzi, G. & Iudicone, D. (2022). Trade-off between sex and growth in diatoms: Molecular mechanisms and demographic implications. Science Advances 8, eabj9466.

4. Apoya-Horton, M.D., Yin, L., Underwood, G.J.C. & Gretz, M.R. (2006). Movement modalities and responses to environmental changes of the mudflat diatom *Cylindrotheca closterium*. Journal of Phycology 42, 379–390.

5. Arora, N. and Philippidis, G.P. (2021). Microalgae strain improvement strategies: Random mutagenesis and adaptive laboratory evolution. Trends in Plant Science 26, pp.1199–1200.

6. Bilcke, G., Campese, L., Annunziata, R., Amadei Martínez, L., Borgonuovo, C., Rijsdijk, N., Chaerle, P., Van den Berge, K., D’hondt, S., Iudicone, D. & Montresor, M. (2025). Conserved genetic markers reveal widespread diatom sexual reproduction in the global ocean. Nature Communications 16, 10029.

7. Belišová, D., Bilcke, G., Audoor, S., D’hondt, S., De Veylder, L., Vandepoele, K. and Vyverman, W. (2025). Molecular fingerprints of cell size sensing and mating type differentiation in pennate diatoms. New Phytologist 245, pp.1625–1639.

8. Bondoc, K.G.V., Lembke, C., Lang, S.N., Germerodt, S., Schuster, S., Vyverman, W. & Pohnert, G. (2019). Decision-making of the benthic diatom *Seminavis robusta* searching for inorganic nutrients and pheromones. The ISME Journal 13, 537–546.

9. Bondoc-Naumovitz, K.G., Crosato, E. & Wan, K.Y. (2025). Functional morphology of gliding motility in benthic diatoms. Proceedings of the National Academy of Sciences USA 122, e2426910122.

10. Camus, M.F., Alexander-Lawrie, B., Sharbrough, J. & Hurst, G.D.D. (2022). Inheritance through the cytoplasm. Heredity 129, 31–43.

11. Casteleyn, G., Adams, N.G., Vanormelingen, P., Debeer, A.E., Sabbe, K. & Vyverman, W. (2009). Natural hybrids in the marine diatom *Pseudo-nitzschia pungens*: genetic and morphological evidence. Protist 160, 343–354.

12. Chepurnov, V.A., Mann, D.G., Von Dassow, P., Vanormelingen, P., Gillard, J., Inzé, D., Sabbe, K. & Vyverman, W. (2008). In search of new tractable diatoms for experimental biology. BioEssays 30, 692–702.

13. Chung, K.P., Gonzalez-Duran, E., Ruf, S., Endries, P. & Bock, R. (2023). Control of plastid inheritance by environmental and genetic factors. Nature Plants 9, 68–80.

14. Chung, K.P. (2025). Cytoplasmic inheritance: The transmission of plastid and mitochondrial genomes across cells and generations. Plant physiology 198, p.kiaf168.

15. Coelho, S.M. & Umen, J.G. (2021). Switching it up: algal insights into sexual transitions. Plant Reproduction 34, 287–296.

16. Collier, S.L., Farrell, S.N., Goodman, C.D. and McFadden, G.I. (2025). Modes and mechanisms for the inheritance of mitochondria and plastids in pathogenic protists. PLoS Pathogens 21, p.e1012835.

17. D’Alelio, D. & Ruggiero, M.V. (2015). Interspecific plastidial recombination in the diatom genus *Pseudo-nitzschia*. Journal of Phycology 51, 1024–1028.

18. Daugbjerg, N. & Andersen, R.A. (1997). A molecular phylogeny of the heterokont algae based on chloroplast-encoded *rbcL* sequence data. Journal of Phycology 33, 1031–1041.

19. De Decker, S., Vanormelingen, P., Pinseel, E., Sefbom, J., Audoor, S., Sabbe, K. & Vyverman, W. (2018). Incomplete reproductive isolation between genetically distinct sympatric clades of the pennate model diatom *Seminavis robusta*. Protist 169, 569–583.

20. Détain, A., Suzuki, H., Wijffels, R.H., Leborgne-Castel, N. and Hulatt, C.J. (2025). Snow algae exhibit diverse motile behaviors and thermal responses. MBio 16, pp.e02954–24.

21. Ferderer, A., Schulz, K.G., Willis, A., Baker, K.G., Chase, Z. and Bach, L.T. (2025). Carbonate chemistry fitness landscapes inform diatom resilience to future perturbations. Science advances 11, p.eadu8024.

22. Glaser, K. & Karsten, U. (2020). Salinity tolerance in biogeographically different strains of the marine benthic diatom *Cylindrotheca closterium*. Journal of Applied Phycology 32, 3809–3816.

23. Golfier, S., Geyer, V.F., Lettermann, L., Schwarz, U.S., Poulsen, N. and Diez, S. (2026). Dynamic switching of cell–substrate contact sites allows gliding diatoms to modulate the curvature of their paths. Proceedings of the National Academy of Sciences 123, p.e2506122123.

24. Goulet, B.E., Roda, F. & Hopkins, R. (2017). Hybridization in plants: old ideas, new techniques. Plant Physiology 173, 65–78.

25. Hofstatter, P.G., Brown, M.W. & Lahr, D.J.G. (2018). Comparative genomics supports sex and meiosis in diverse Amoebozoa. Genome Biology and Evolution 10, 3118–3128.

26. Huang, Y., Sheth, R.U., Zhao, S., Cohen, L.A., Dabaghi, K., Moody, T., Sun, Y., Ricaurte, D., Richardson, M., Velez-Cortes, F. & Blazejewski, T. (2023). High-throughput microbial culturomics using automation and machine learning. Nature Biotechnology 41, 1424–1433.

27. Jankowicz-Cieslak, J., Mba, C. and Till, B.J. (2016). Mutagenesis for crop breeding and functional genomics. In Biotechnologies for plant mutation breeding: protocols (pp. 3–18). Cham: Springer International Publishing.

28. Klapper, F., Audoor, S., Vyverman, W. & Pohnert, G. (2021). Pheromone-mediated sexual reproduction of the pennate diatom *Cylindrotheca closterium*. Journal of Chemical Ecology 47, 504–512.

29. Krishnan, A., Cano, M., Karns, D.A., Burch, T.A., Likhogrud, M., Aqui, M., Bailey, S., Verruto, J., Lambert, W., Kuzminov, F. and Naghipor, M. (2023). Simultaneous CAS9 editing of cpSRP43, LHCA6, and LHCA7 in Picochlorum celeri lowers chlorophyll levels and improves biomass productivity. Plant Direct 7, p.e530.

30. Krueger-Hadfield, S.A. (2024). Let’s talk about sex: why reproductive systems matter for understanding algae. Journal of Phycology 60, 581–597.

31. LaPanse, A.J., Krishnan, A. & Posewitz, M.C. (2021). Adaptive laboratory evolution for algal strain improvement: methodologies and applications. Algal Research 53, 102122.

32. LaPanse, A.J., Burch, T.A., Tamburro, J.M., Traller, J.C., Pinowska, A. and Posewitz, M.C. (2023). Adaptive laboratory evolution for increased temperature tolerance of the diatom Nitzschia inconspicua. MicrobiologyOpen 12, p.e1343.

33. Li, Z., Zhang, Y., Irwin, A.J. and Finkel, Z.V. (2025). Polyploidization in diatoms accelerates adaptation to warming. Nature Climate Change 15, 1241–1248.

34. Mann, D.G. (1999). The species concept in diatoms. Phycologia 38, 437–495.

35. Mann, D.G. & Vanormelingen, P. (2013). An inordinate fondness? The number, distributions, and origins of diatom species. Journal of Eukaryotic Microbiology 60, 414–420.

36. Marshall, W.F., Young, K.D., Swaffer, M., Wood, E., Nurse, P., Kimura, A., Frankel, J., Wallingford, J., Walbot, V., Qu, X. & Roeder, A.H.K. (2012). What determines cell size? BMC Biology 10, 101.

37. Monnet, F., Postel, Z., Touzet, P., Fraïsse, C., Van de Peer, Y., Vekemans, X. & Roux, C. (2025). Rapid establishment of species barriers in plants compared with that in animals. Science 389, 1147–1150.

38. Montresor, M., Vitale, L., D’Alelio, D. & Ferrante, M.I. (2016). Sex in marine planktonic diatoms: insights and challenges. Perspectives in Phycology 3, 61–75.

39. Moore, E.R., Bullington, B.S., Weisberg, A.J., Jiang, Y., Chang, J. & Halsey, K.H. (2017). Morphological and transcriptomic evidence for ammonium induction of sexual reproduction in *Thalassiosira pseudonana* and other centric diatoms. PLoS ONE 12, e0181098.

40. Naseeb, S., Visinoni, F., Hu, Y., Hinks-Roberts, A.J., Maslowska, A., Walsh, T., Smart, K.A., Louis, E.J. & Delneri, D. (2021). Restoring fertility in yeast hybrids: breeding and quantitative genetics of beneficial traits. PNAS 118, e2101242118.

41. Oliver, A., Podell, S., Pinowska, A., Traller, J.C., Smith, S.R., McClure, R., Beliaev, A., Bohutskyi, P., Hill, E.A., Rabines, A. and Zheng, H. (2021). Diploid genomic architecture of Nitzschia inconspicua, an elite biomass production diatom. Scientific Reports 11, p.15592.

42. Prigent, L., Quéré, J., Plus, M. & Le Gac, M. (2025). Sexual reproduction during diatom blooms. ISME Communications 5, ycae169.

43. Ruggiero, M.V., Buffoli, M., Wolf, K.K., D’Alelio, D., Di Tuccio, V., Lombardi, E., Manfellotto, F., Vitale, L., Margiotta, F., Sarno, D. & John, U. (2024). Multiannual patterns of genetic structure and mating-type ratios highlight complex bloom dynamics of a marine planktonic diatom. Scientific Reports 14, 6028.

44. Speijer, D., Lukeš, J. & Eliáš, M. (2015). Sex is a ubiquitous, ancient, and inherent attribute of eukaryotic life. PNAS 112, 8827–8834.

45. Stock, W., Vanelslander, B., Rüdiger, F., Sabbe, K., Vyverman, W. & Karsten, U. (2019). Thermal niche differentiation in the benthic diatom *Cylindrotheca closterium* complex. Frontiers in Microbiology 10, 1395.

46. Tanaka, T., Maeda, Y., Veluchamy, A., Tanaka, M., Abida, H., Maréchal, E., Bowler, C., Muto, M., Sunaga, Y., Tanaka, M. and Yoshino, T. (2015). Oil accumulation by the oleaginous diatom Fistulifera solaris as revealed by the genome and transcriptome. The Plant Cell 27, pp.162–176.

47. Vanormelingen, P., Vanelslander, B., Sato, S., Gillard, J., Trobajo, R., Sabbe, K. & Vyverman, W. (2013). Heterothallic sexual reproduction in the model diatom *Cylindrotheca closterium*. European Journal of Phycology 48, 93–105.

48. Yadav, S., Ghosh, D., Twinkle & Dogra, V. (2025). Biparental plastid inheritance and its implications in plastid genome engineering. Plant Cell Reports 44, 236.

49. Zhang, Q., Leng, H.T., Li, H., Arrigo, K.R. & Prakash, M. (2025). Ice-gliding diatoms establish record-low temperature limits for motility in a eukaryotic cell. Proceedings of the National Academy of Sciences USA 122, e2423725122.

50. Zheng, B., Lucas, A.J., Franks, P.J.S., Schlosser, T.L., Anderson, C.R., Send, U., Davis, K., Barton, A.D. & Sosik, H.M. (2023). Dinoflagellate vertical migration fuels an intense red tide. PNAS 120, e2304590120.

