## Supporting data for "Experimental breeding reveals mating patterns and F1 phenotypic diversification in the cosmopolitan diatom *Cylindrotheca*"

### Supporting data S1: Examples of mating cells

Mating *cyr*1 x *cyr*2 small morphotype (Clade C) ×400 well plate

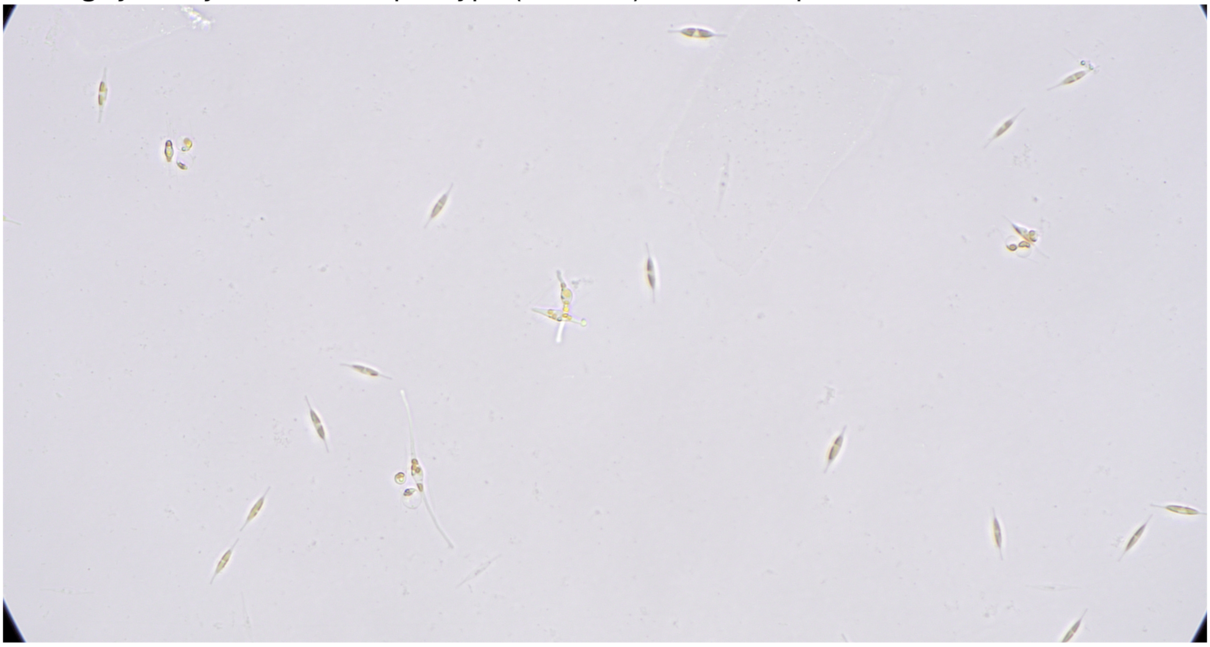

Mating *cyr*24 x *cyr*38 large morphotype (Clade B) ×200 well plate

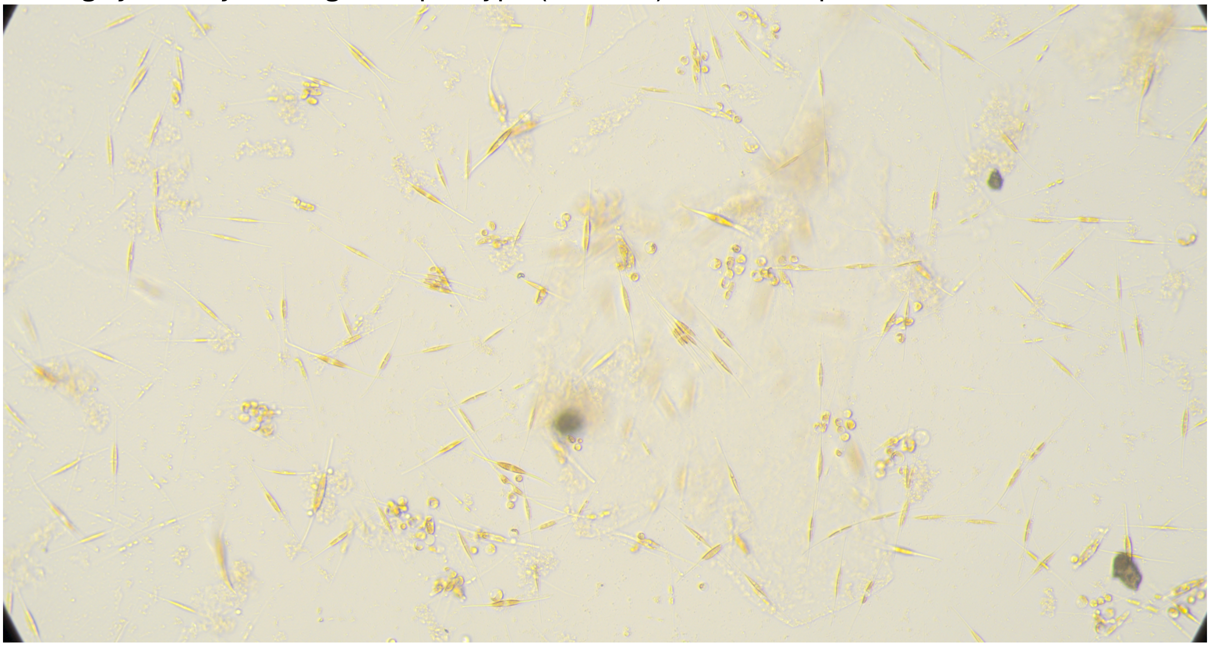

**Supporting data S2: PCA analysis**

Scree plot

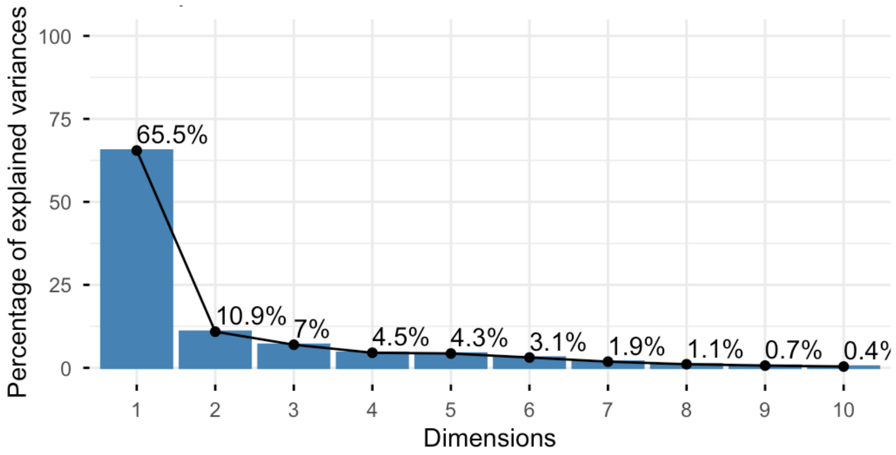


PCA loadings

| | PC1 | PC2 | PC3 |
|---|---|---|---|
| **growth** | -0.07 | 0.52 | -0.63 |
| **length** | 0.32 | -0.01 | -0.13 |
| **area** | 0.31 | -0.16 | -0.16 |
| **chlorophyll** | 0.27 | -0.08 | -0.39 |
| **yield_blue** | 0.15 | 0.56 | 0.06 |
| **yield_green** | 0.23 | 0.4 | 0.31 |
| **alpha_blue** | 0.27 | 0.17 | -0.09 |
| **alpha_green** | 0.27 | 0.15 | -0.04 |
| **etr_blue** | 0.28 | 0.15 | 0.24 |
| **etr_green** | 0.26 | 0.11 | 0.43 |
| **mean_speed** | 0.31 | -0.15 | -0.07 |
| **max_speed** | 0.3 | -0.19 | 0.04 |
| **confinement** | 0.29 | -0.22 | -0.22 |
| **direction** | -0.29 | 0.16 | 0 |

**Supporting data S3: Plastid genome inheritance in four F1 *Cylindrotheca* offspring.**

Sequence reads from 4 different offspring were mapped to the *cyr*1 parental *rbc*L haplotype. Highly accurate HiFi reads at QV34-35 were mapped in Geneious Prime with minimap2 for PacBio reads. The following are the alignment graph and variants from the four F1 sequence read datasets, and a result summary.

| Cross | F1 isolate | Parent plastid origin |
|---|---|---|
| *cyr*1 x *cyr*2 | A | *cyr*2 only |
| *cyr*1 x *cyr*2 | B | *cyr*2 only |
| *cyr*1 x *cyr*4 | A | *cyr1* only |
| *cyr*1 x *cyr*4 | G | *cyr4* only |

**F1 cross between *cyr*1 x *cyr*2, isolate A**. Has 6 SNPs corresponding to *cyr*2 and has inherited only the ***cyr*2 parent plastid genome** and **no evidence of heteroplasmy.**

| Minimum | Maximum | Length | Change | Coverage | Polymorphism Type | Strand-Bias | Variant Frequency | Variant P-Value (approxim... |
|---|---|---|---|---|---|---|---|---|
| 1,190 | 1,190 | 1 | C -> T | 574 | SNP (transition) | 53.8% | 99.5% | 0.0 |
| 1,026 | 1,026 | 1 | G -> A | 570 | SNP (transition) | 53.3% | 100.0% | 0.0 |
| 1,042 | 1,042 | 1 | A -> G | 571 | SNP (transition) | 53.3% | 99.3% | 0.0 |
| 365 | 365 | 1 | A -> G | 566 | SNP (transition) | 52.6% | 96.8% | 0.0 |
| 356 | 356 | 1 | C -> T | 566 | SNP (transition) | 52.3% | 100.0% | 0.0 |
| 116 | 116 | 1 | G -> A | 557 | SNP (transition) | 52.1% | 99.6% | 0.0 |

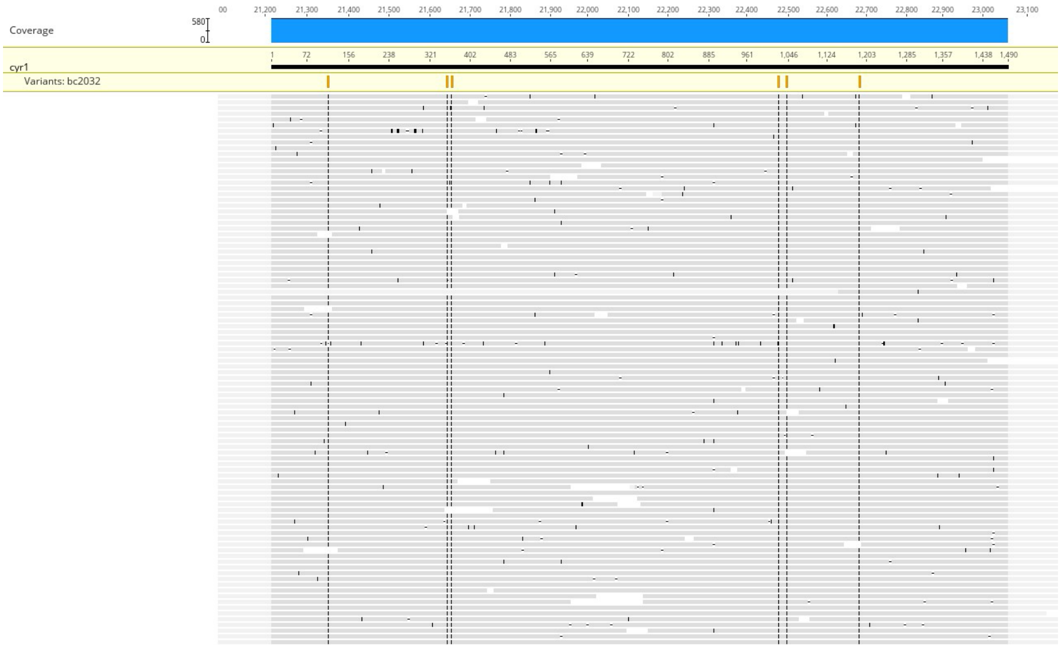


**F1 cross between *cyr*1 x *cyr*2, isolate B**. Has 6 SNPs corresponding to *cyr*2 and has inherited only the ***cyr*2 parent plastid genome** and **no evidence of heteroplasmy.**

| Minimum | Maximum | Length | Change | Coverage | Polymorphism Type | Strand-Bias | Variant Frequency | Variant P-Value (approxim... |
|---|---|---|---|---|---|---|---|---|
| 356 | 356 | 1 | C -> T | 662 | SNP (transition) | 53.3% | 100.0% | 0.0 |
| 116 | 116 | 1 | G -> A | 657 | SNP (transition) | 53.2% | 99.2% | 0.0 |
| 1,190 | 1,190 | 1 | C -> T | 672 | SNP (transition) | 53.2% | 99.1% | 0.0 |
| 365 | 365 | 1 | A -> G | 661 | SNP (transition) | 53.1% | 97.7% | 0.0 |
| 1,042 | 1,042 | 1 | A -> G | 671 | SNP (transition) | 53.1% | 99.6% | 0.0 |
| 1,026 | 1,026 | 1 | G -> A | 673 | SNP (transition) | 53.0% | 100.0% | 0.0 |

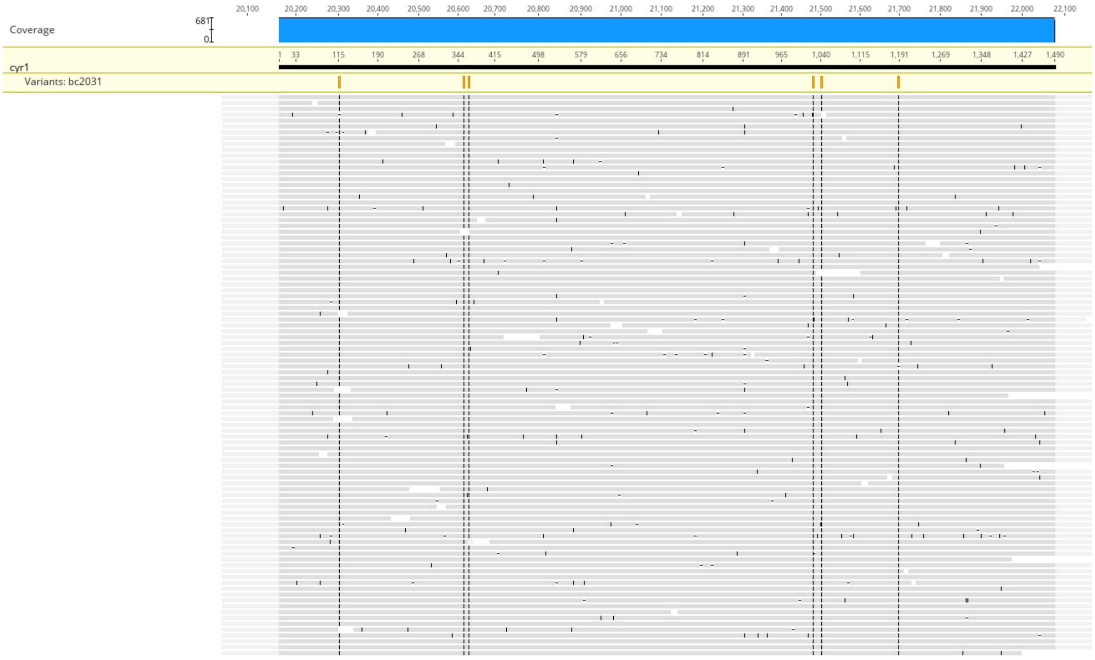


**F1 cross between *cyr*1 x *cyr*4, isolate A**. Identical to *cyr*1 and has inherited only the ***cyr*1 parent plastid genome** and **no evidence of heteroplasmy.**

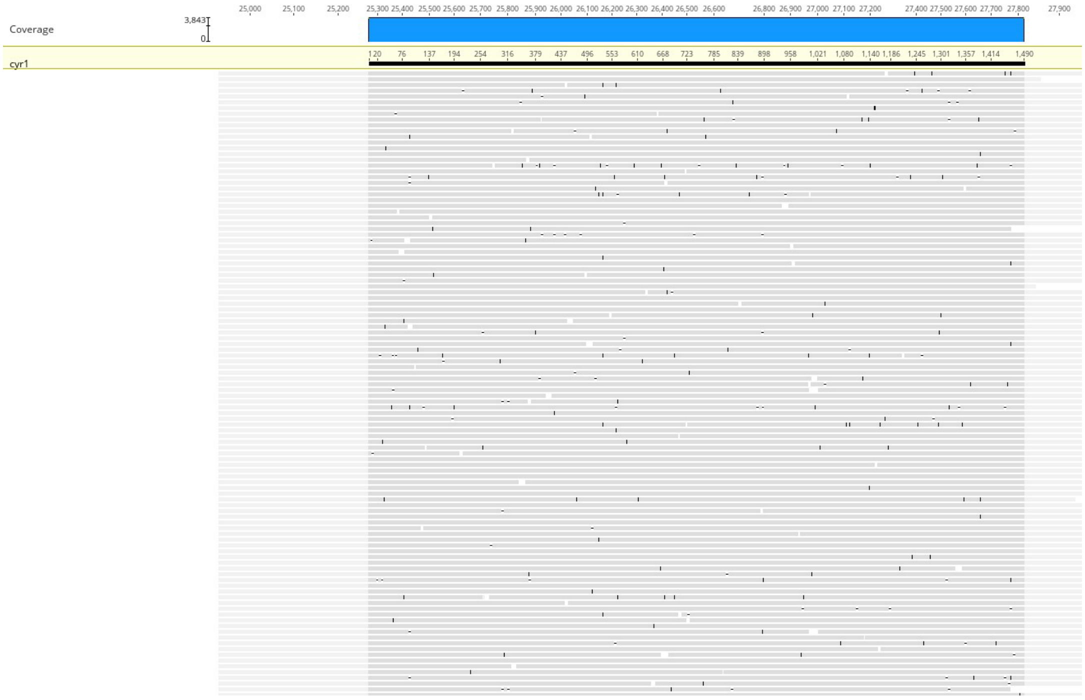

**F1 cross between *cyr*1 x *cyr*4, isolate G**. Has 6 SNPs corresponding to *cyr*4 and has inherited only the ***cyr*4 parent plastid genome** and **no evidence of heteroplasmy.**

| Minimum | Maximum | Length | Change | Coverage | Polymorphism Type | Strand-Bias | Variant Frequency | Variant P-Value (approxim... |
|---|---|---|---|---|---|---|---|---|
| 365 | 365 | 1 | A -> G | 7,231 | SNP (transition) | 50.8% | 97.1% | 0.0 |
| 116 | 116 | 1 | G -> A | 7,215 | SNP (transition) | 50.7% | 99.5% | 0.0 |
| 356 | 356 | 1 | C -> T | 7,229 | SNP (transition) | 50.7% | 99.8% | 0.0 |
| 1,190 | 1,190 | 1 | C -> T | 7,325 | SNP (transition) | 50.6% | 99.6% | 0.0 |
| 1,026 | 1,026 | 1 | G -> A | 7,286 | SNP (transition) | 50.5% | 100.0% | 0.0 |
| 1,042 | 1,042 | 1 | A -> G | 7,288 | SNP (transition) | 50.5% | 99.3% | 0.0 |

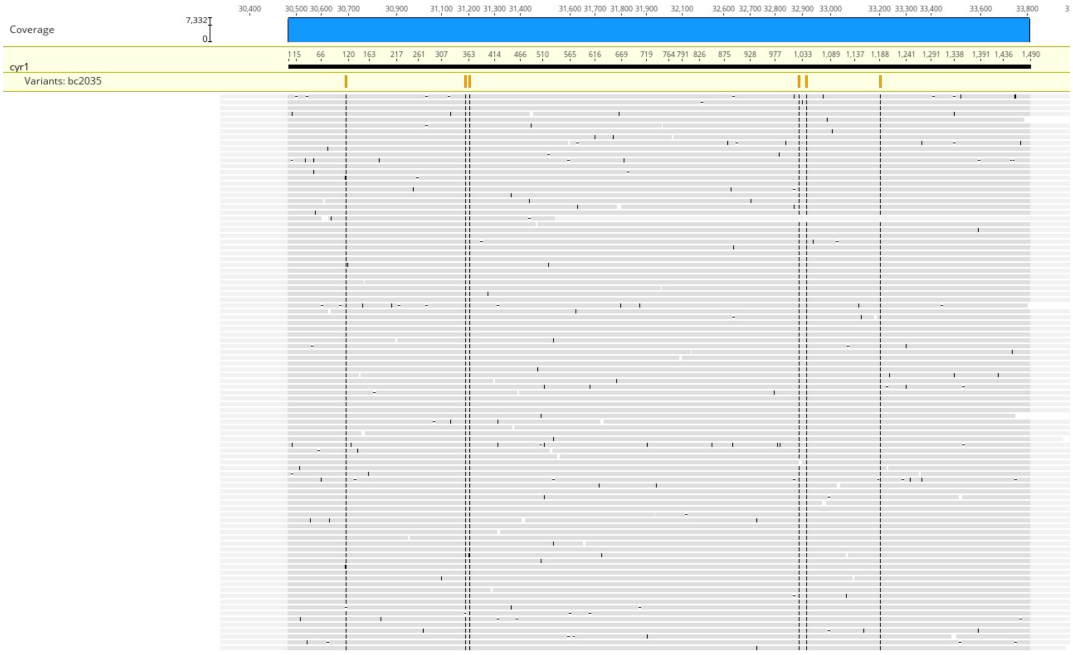
